# Flawed evidence for convergent evolution of the circadian *CLOCK* gene in mole-rats

**DOI:** 10.1101/022004

**Authors:** Frédéric Delsuc

**Author notes:** ARISING FROM X. Fang *et al. Nature Communications* 5, 3966 (2014); doi:10.1038/ncomms4966.

## Abstract

Convergently evolved mole-rats (Mammalia, Rodentia) provide a fascinating model for studying convergent molecular evolution. Three genome sequences have recently been made available for the blind mole-rat (*Nannospalax galili*; Spalacidae; Muroidea)^1^, and the convergently evolved naked mole-rat (*Heterocephalus glaber*; Heterocephalidae; Ctenohystrica)^2^ and its close relative the Damaraland mole-rat (*Fukomys damarensis*; Bathyergidae; Ctenohystrica)^3^. In their genome paper^1^, Fang *et al.* evaluated convergent molecular evolution related to the subterranean life-style between the naked mole-rat and the blind mole-rat. One particularly striking result was the strong signal for amino acid convergence detected in the circadian rhythm *CLOCK* gene. Here I show that this unexpected result is erroneous because it is based on the use of the wrong sequence for the naked mole-rat, which has been mistakenly replaced by a sequence from a blind mole-rat. When the correct sequence is used, the evidence for convergent molecular evolution in this gene appears very limited.

---

In their paper reporting the genome of the blind mole-rat**^1^**, Fang et al. studied convergent evolution related to underground stresses between *Nannospalax galili* and *Heterocephalus glaber*. As reported in their Figure 1, these two mole-rat species evolved convergently in two distinct rodent clades that are estimated to have diverged more than 70 million years ago. In this context, one particularly striking result was the astonishingly high degree of amino acid convergence found in the *CLOCK* gene, which is involved in the regulation of the circadian rhythm. The evidence for convergent molecular evolution stems from an alignment of the last 173 sites of the CLOCK protein in four rodent species (*Mus*, *Rattus*, *Heterocephalus*, and *Nannospalax*) and human (*Homo*) presented in their Figure 2a. Fang et al.**^1^** observed that, based on these 173 amino acid sites, the CLOCK proteins of *Heterocephalus* and *Nannospalax* are identical, including in a Glutamine-rich region believed to function as a tuning knob regulating circadian rhythmicity**^4^**. The phylogenetic tree of the CLOCK protein they presented as Figure 2b accordingly groups the two convergently evolved mole-rat species. The strong bootstrap support obtained for their monophyly, and the very short terminal branches exhibited by these two species imply that their complete CLOCK protein sequences should be almost identical. Based on these observations, Fang et al.**^1^** concluded that the *CLOCK* gene has been strongly shaped by convergent molecular evolution in these independently evolved subterranean animals.

**Figure 1.**
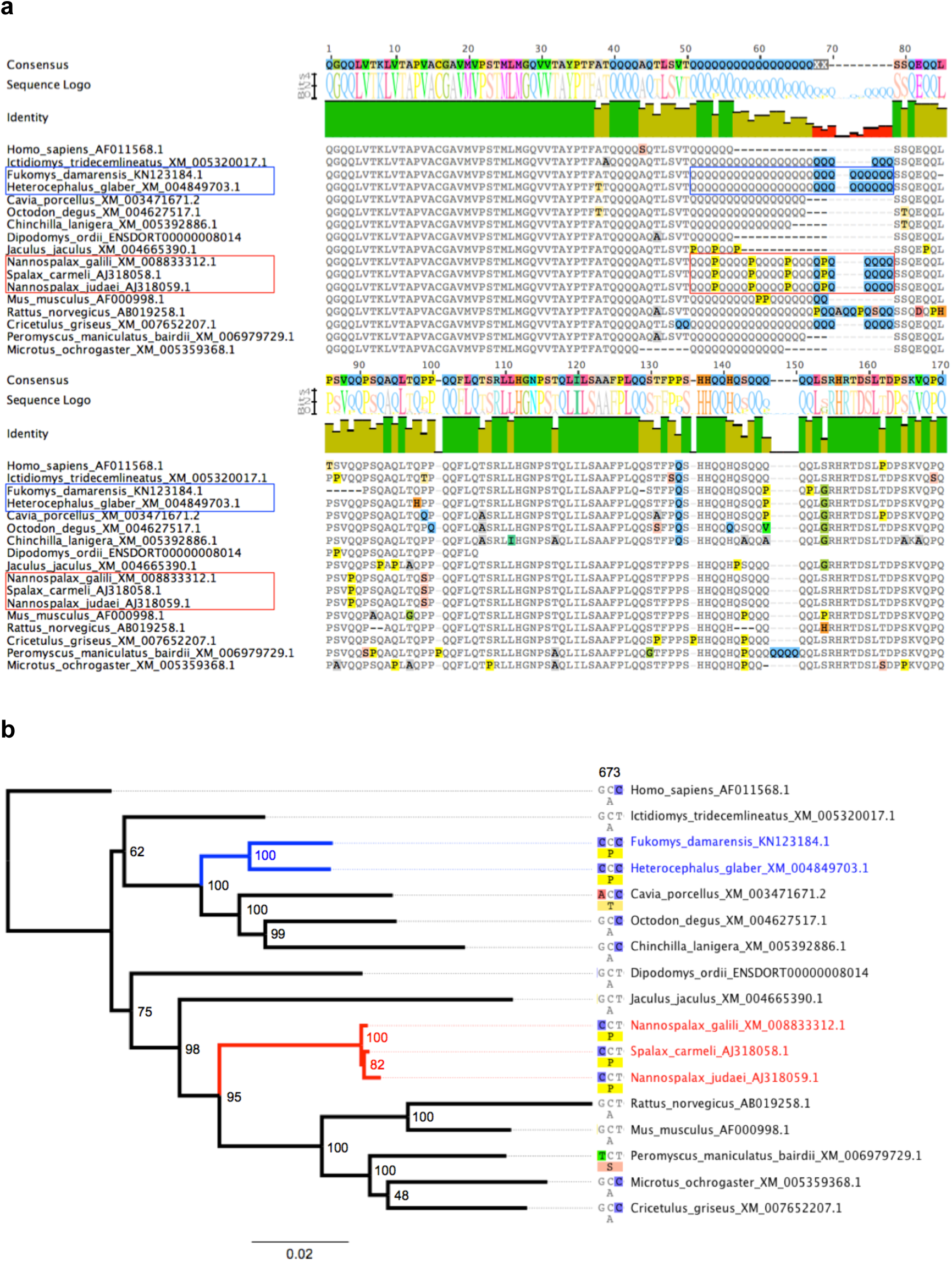
Corrected alignment and phylogeny of the CLOCK gene. (**a**) Alignment of the last 170 amino acid sites for all 16 available rodent CLOCK protein sequences plus *Homo sapiens*. The Glutamine-rich region is boxed in the two convergent mole-rat groups (Bathyergidae and Heterocephalidae in blue, and Spalacidae in red). (**b**) Maximum likelihood phylogenetic tree inferred from the corresponding codon alignment of the *CLOCK* gene. Numbers at nodes indicate bootstrap percentages. The single potentially convergent site detected (amino acid position 673) is shown.

This result is particularly surprising since convergent evolution is expected to typically affect only a limited number of positions with functional effect in a given protein**^5,6^**. Convergent molecular evolution is certainly not expected to result in two protein sequences with such a high level of similarity in two distantly related species. Despite the extensive supplementary material associated with Fang et al.’s paper**^1^**, the only information on the sequences and methodology used to infer the alignment and phylogenetic tree is restricted to the legend of their Figure 2, which is unfortunately rather uninformative. It is thus impossible to know which phylogenetic reconstruction method and sequences have been used, all the more so that the taxon sampling differs between the alignment of their Figure 1a and the tree of their Figure 2b. Moreover, the CLOCK sequence for *Spermophilus dauricus* included in the tree of Figure 2b is not available in public databases.

In order to verify this suspicious result, I collected and aligned all 16 available rodent *CLOCK* sequences and reconstructed their phylogenetic relationships using maximum likelihood. As shown in Fig. 1a, the new alignment revealed a number of amino acid changes between the convergent mole-rat groups. In fact, on the last 170 sites of the alignment, all three spalacid sequences (including *Nannospalax*) are identical. However, they strongly differ from the sequences of the convergently evolved *Heterocephalus* and *Fukomys*. In particular, the glutamine-rich region is regularly interrupted by Proline substitutions in spalacids whereas it is composed of only poly-Glutamines in the naked and Damaraland mole-rat sequences. Based on this partial alignment, *Heterocephalus* and the three identical spalacid sequences differ by 13 amino acids. Overall, there are actually 39 amino acid differences between the *Heterocephalus* and *Nannospalax* CLOCK protein sequences (Supplementary Figure S1).

The phylogenetic tree obtained from the complete codon alignment of these CLOCK sequences (Fig. 1b) is fully compatible with the current understanding of rodent phylogeny**^7^** in strongly supporting the independent origins of the two mole-rat groups. This topology strongly conflicts with the one obtained by Fang et al.**^1^** for the CLOCK protein, which supported the monophyly of *Heterocephalus* and *Nannospalax*, and was interpreted as strong support for convergent evolution as a consequence of the subterranean lifestyle. The most likely explanation for this conflicting result is that the *Heterocephalus CLOCK* sequence has been erroneously replaced by a spalacid sequence in Fang et al.’s alignment. The phylogenetic tree of available *CLOCK* sequences (Fig. 1b) does not support the hypothesis of convergent molecular evolution. In fact, screening the amino acid alignment for molecular convergence detects only one potentially convergent site at position 673 on a total of 860 sites. This site presents a convergent amino acid change (Adenine -> Proline) in the two independent mole-rat lineages caused by parallel substitutions leading to two different but synonymous codons (Fig. 2b). This particular site might possibly have a functional effect on the protein activity, however in the absence of any specific functional assays, the evidence for convergent molecular evolution of the CLOCK protein between mole-rats currently appears rather weak, if not non-existent.

## Methods

Complete CLOCK coding sequences (CDSs) were extracted from GenBank and from mammalian genome assemblies available at NCBI for 16 rodent species and *Homo sapiens* used as an outgroup. These coding sequences have been aligned using MACSE**^8^**, which returns both codon and amino acid alignments. Maximum likelihood phylogenetic reconstruction was conducted on the codon alignment using the PhyML 3.0 plugin**^9^** of Geneious R7**^10^** under the GTR+G8 model using SPR branch swapping on a BioNJ starting tree with 100 booststrap replications. Detection of convergent amino acid sites in the CLOCK protein was performed by comparing site-wise likelihood differences**^6^** as estimated by PhyML between the ML topology and two alternative convergent topologies where mole-rats have been constrained to be monophyletic either within Ctenohystrica or within the mouse-related clade.

